## Supplementary Figures for "Human centromere formation activates transcription and opens chromatin fibre structure"

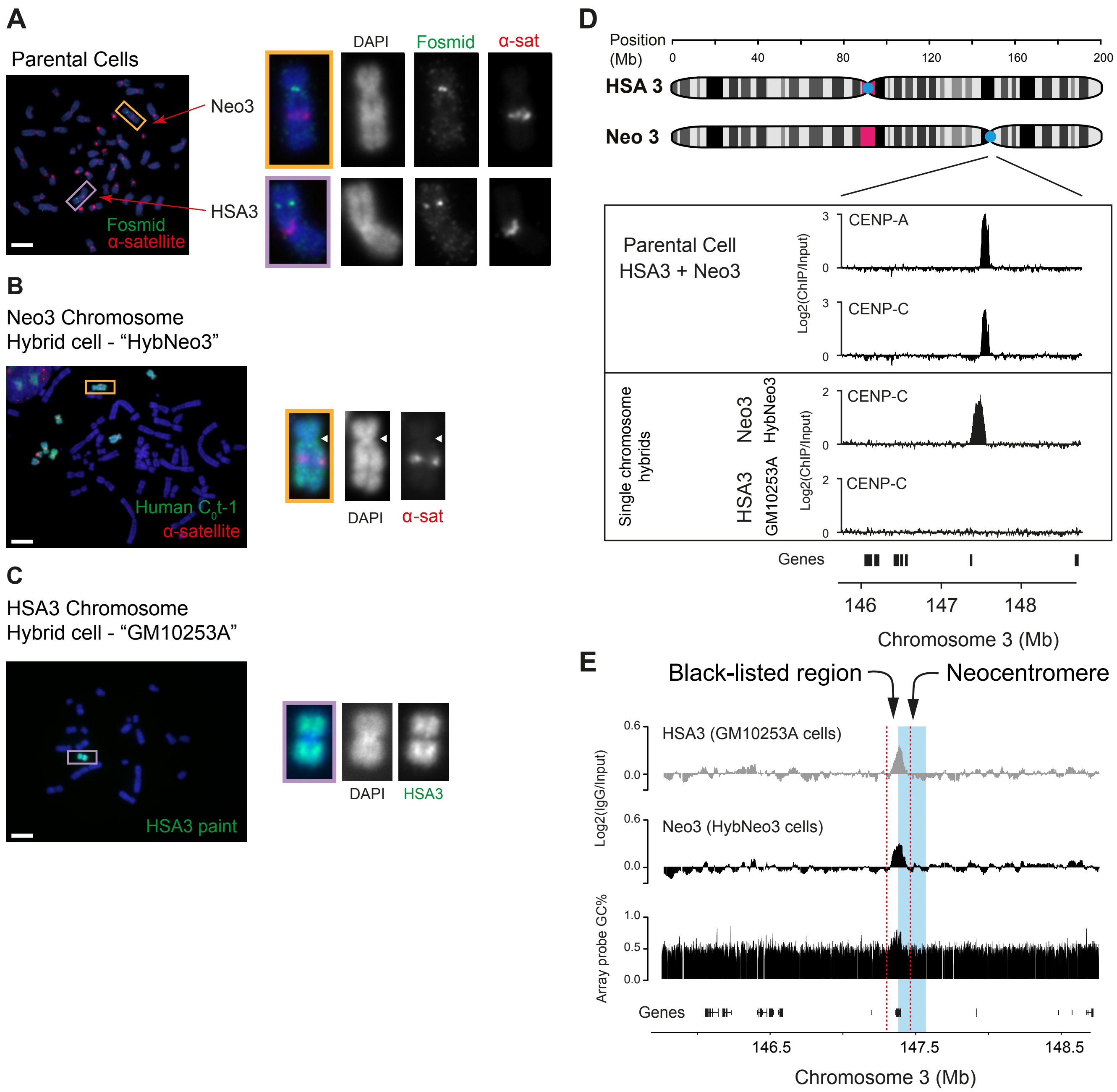


**Supplementary Fig. 1. Cell line model system used to interrogate centromeric chromatin structure and mapping of neocentromere mapping to 3q24 (relates to Figure 1).**
**(A)** Parental human lymphoblastoid cells with one canonical chromosome 3 (HSA3, purple frame) and one chromosome 3 with the centromere relocated to form a neocentromere at 3q24 (Neo3, orange frame). DNA FISH with a-satellite specific probe (red) and a 3q24 fosmid probe (green). **(B)** The Neo3 chromosome was retained after fusion of the parental cell line with a hamster cell to create a hybrid line called “HybNeo3”. Metaphase spread of HybNeo3 hybridized with human $C_0$t-1 DNA (green) identifying the seven human chromosomes present in this human/hamster hybrid (4,6,8,11,13,18, X and Neo3 (orange frame)) and a human a-satellite specific FISH probe (red). **(C)** The GM10253A hybrid cell line has a single canonical human chromosome 3 (HSA3). Metaphase spread of GM10253A hybridized with a human chromosome 3 paint (green; purple frame). Nuclei are counterstained with DAPI. Bar is 5 µm. **(D)** Top, ideogram depicting chromosomes HSA3 and Neo3. Bottom, distribution of CENP-A and CENP-C ChIP signal in parental and hybrid cells at 3q24. **(E)** Top, signal for control IgG ChIP-chip and bottom microarray probe GC composition (%). Blacklisted region (chr3: 147324413-147482213; hg38) is marked by red dashed lines. Neocentromere core (chr3: 147400413-147591023) defined from CENP-C (panel A) is marked in blue.

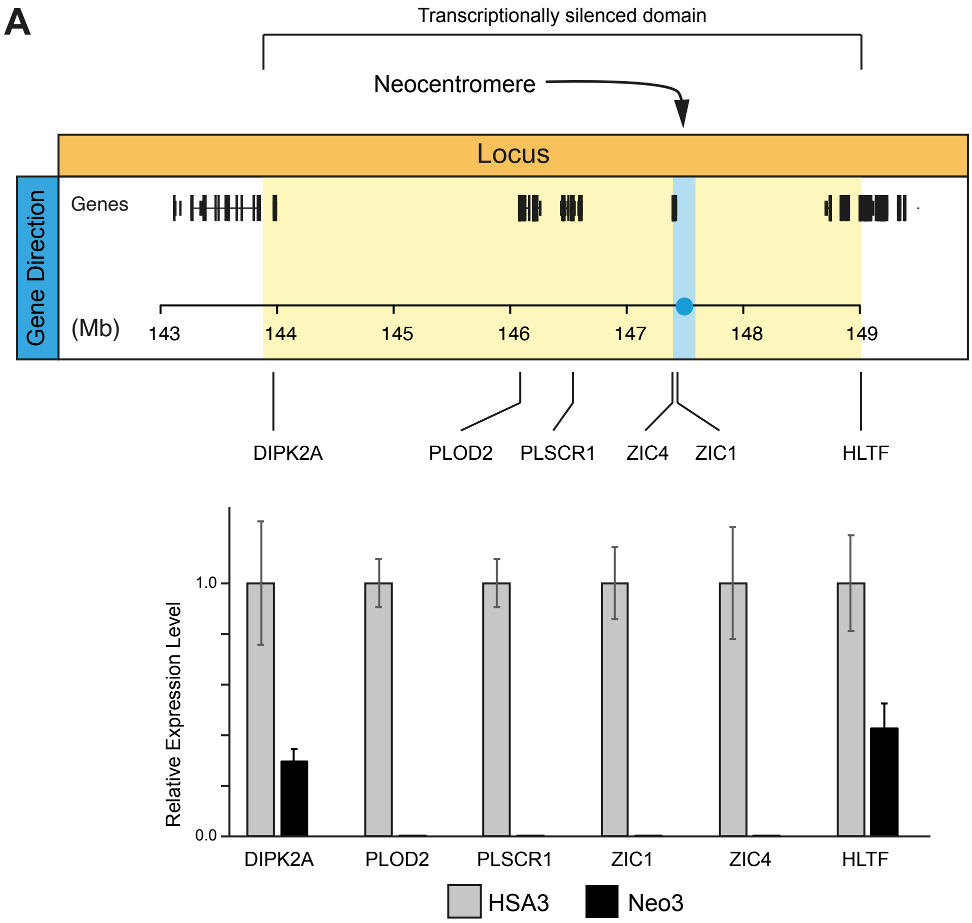


**Supplementary Fig. 2. Transcription repression at the pericentromeric domain (Relates to Figure 2)** Top, diagram showing individual genes at 3q24, yellow block corresponds to pericentromere domain, blue is the neocentromere. Bottom, RT-qPCR expression data for genes within and bordering the pericentromeric heterochromatic domain, showing expression from HSA3 and gene silencing on Neo3.

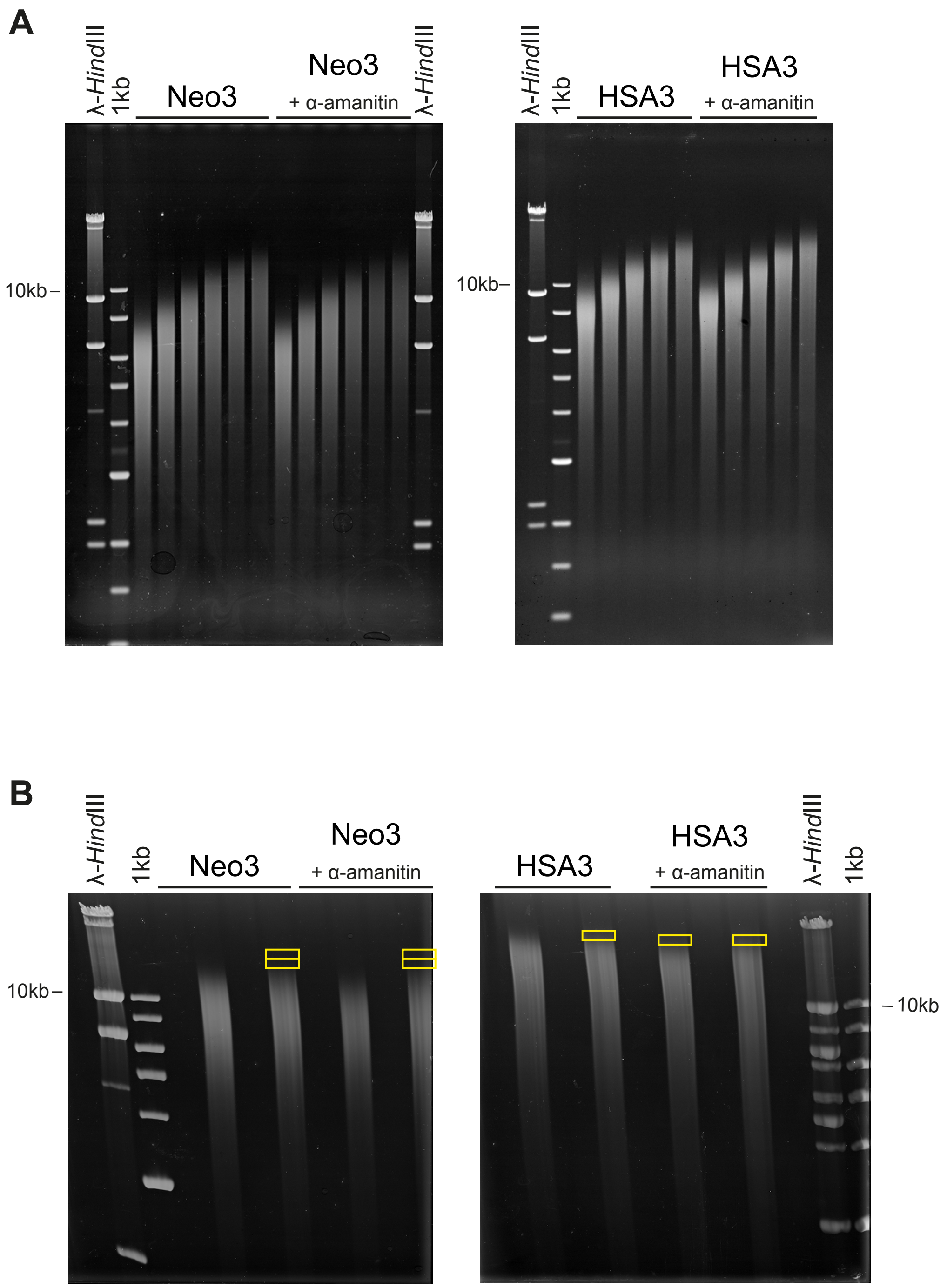


**Supplemenary Fig. 3. Isolation of "open" chromatin probes (Relates to Figure 4).**
Soluble chromatin isolated from hybrid cell nuclei by micrococcal digestion was fractionated by size and structure on a sucrose gradient. **(A)** Top, agarose gel electrophoresis of DNA purified from sucrose gradient fractions. **(B)** DNA from gradient fractions was size selected by PFGE. DNA fragments ≈10 kb longer than the bulk of the DNA signal, corresponding to "open" or disrupted chromatin, was purified from gel slices (yellow boxes).

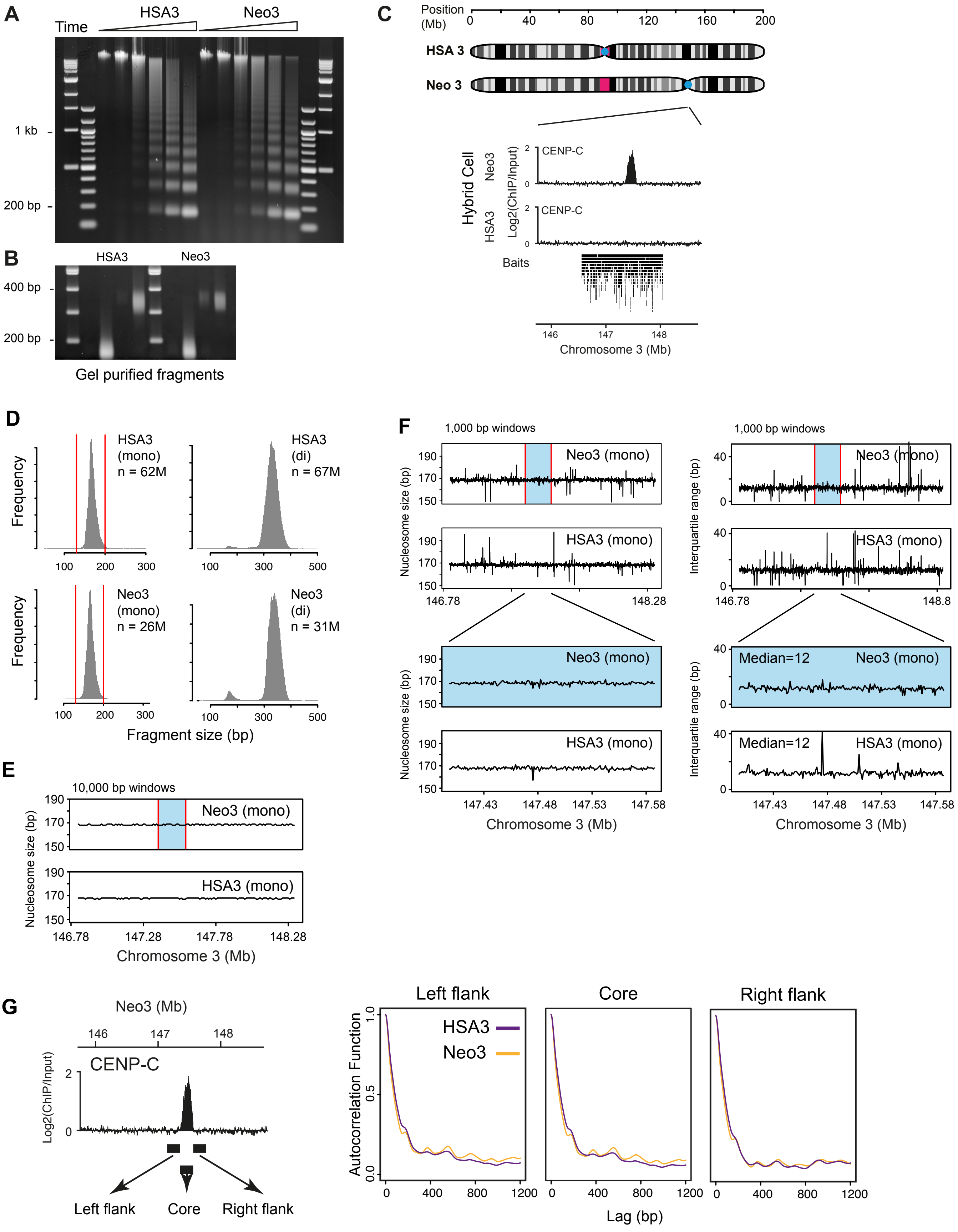


Supplementary Fig. 4
Legend over page

**Supplementary Fig. 4. Arrangement of nucleosomes around neocentromere (Relates to Figure 5).** **(A)** Agarose gel electrophoresis of DFF digested nuclei isolated from HSA3 and Neo3 containing cells. Mono and di-nucleosome fragments were excised, and the DNA extracted using β-agarase. **(B)** Agarose gel electrophoresis of purified mono- and di-nucleosomes fragments used for nucleosome mapping. **(C)** Top, ideogram depicting HSA3 and Neo3 chromosomes with enlargement of 3 Mb region around the neocentromere (marked by CENP-C). Bottom, genomic location of the capture baits used to enrich for 1.5 Mb of neocentromeric region. **(D)** Size distribution of mono and di nucleosomes (bp) isolated from HSA3 and Neo3 cells for 1.5 Mb around the region corresponding to the neocentromere. **(E)** Mono nucleosome size distribution in 10 kb windows across the 1.5 Mb captured domain (neocentromere marked in blue). **(F)** Left, mono nucleosome size (median) in 1 kb windows across the 1.5 Mb captured domain and focussed region covering the neocentromere (marked in blue). Right, variance (interquartile range) in mono nucleosome size in 1 kb windows across the 1.5 Mb captured domain and focussed region covering the neocentromere (marked in blue). **(G)** Autocorrelation of nucleosome dyad coverage at left flank, centromere core and right flank for different lag (bp).

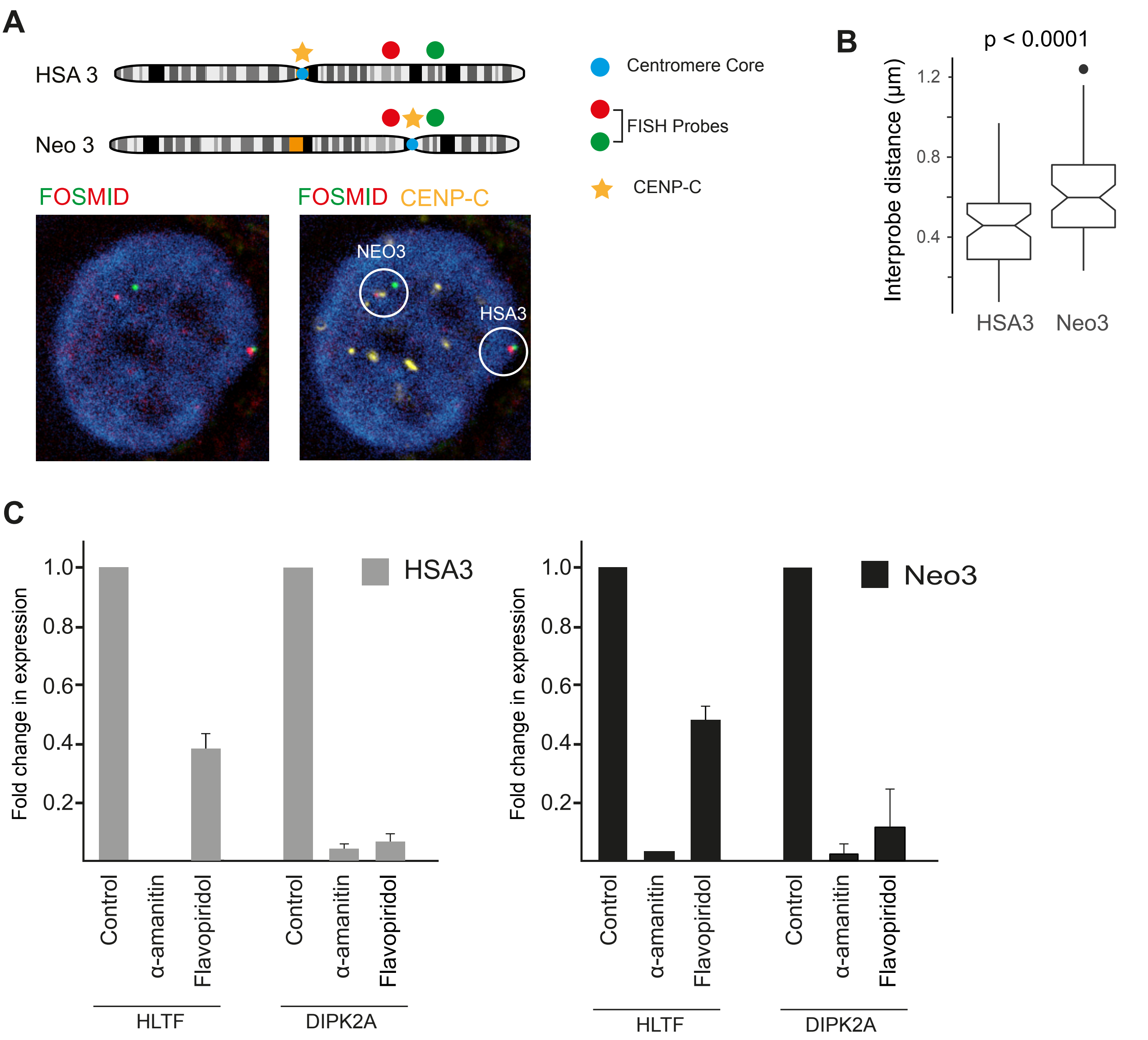


**Supplementary Fig. 5. Large scale chromatin is decompacted at the neocentromere in the parental cell line (Relates to Figure 6).**

**(A)** Top, chromosome 3 ideogram indicating CENP-C immunofluoresence signal (yellow) and the neocentromere specific FISH probes (green and red). Bottom right, representative image of 4 colour 3D immuno-FISH for identifying the chromosome 3 harbouring a neocentromere at 3q24 due to the presence of the CENP-C signal proximal to one pair of fosmid probes. Below left, three colour representation of the same image used for measuring interprobe distance. **(B)** Boxplot showing interprobe distance measurements (µm) between the pair of fosmid probes (B and C, see Fig 6A) for the HSA3 and Neo3 chromosomes in the parental lymphoblastoid cells. **(C)** RT-qPCR expression data for genes in the pericentromere region flanking the neocentromere domain in cells carrying the HSA3 and Neo3 chromosomes, following transcription inhibition with a-amanitin (5h) or flavopiridol (3h).

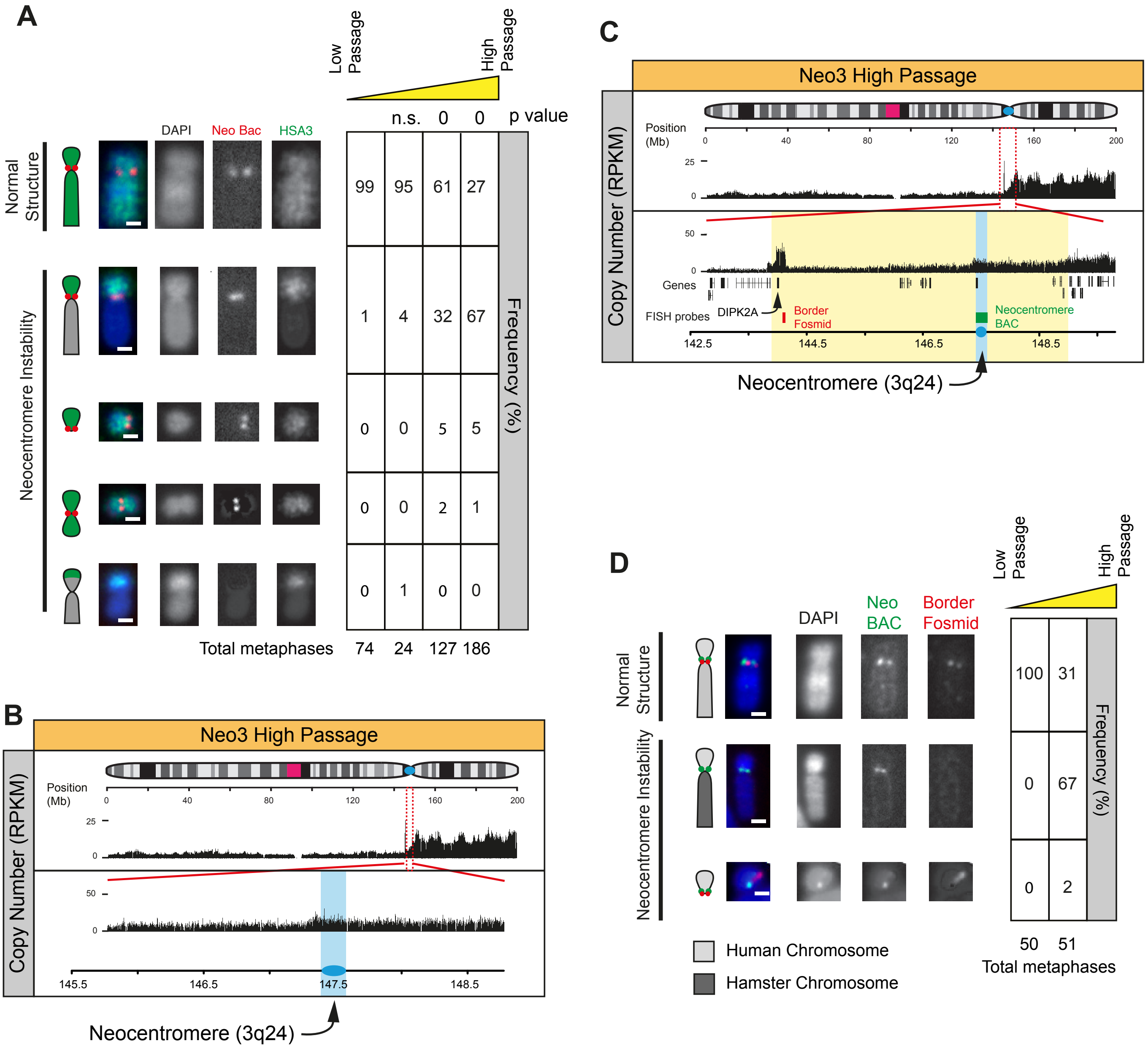


**Supplementary Fig. 6. Neocentromere associated genome instability (Relates to Figure 7)**
**(A)** Left, representative FISH images of Neo3 metaphase chromosomes hybridised to a human chromosome 3 paint (green) and BAC (red) located at the neocentromere. Chromosome morphology was scored as normal or showing instability: deletions, fusions or duplications. Bar is 2μm. Right, quantification (%) of different chromosome morphologies with increasing passage number (low ~ 10, high ~ 100) over time. P values are for a $\chi^2$ test compared to low passage. **(B)** High passage Neo3 Chromosome copy number (RPKM), with a zoom in of the 3 Mb region around the neocentromere (blue). **(C)** High passage Neo3 Chromosome copy number (RPKM), with a zoom in of the 7 Mb region around the neocentromere (blue). Locations of DNA FISH probes are shown, neocentromere BAC probe (green) and border fosmid (red). **(D)** Left, representative FISH images of Neo3 metaphase chromosomes hybridised to a BAC (green) located at the neocentromere and a fosmid (red) located at the border. Bar is 2μm. Right, chromosome morphology was scored and quantified (%) with increasing passage number over time. Loss of the border probe signal was coincident with fusion of human Neo3 fragment (light grey) to a hamster chromosome (dark grey). Bar is 2μm.
